# Confined filaments in soft vesicles - case of sickle red blood cells

**DOI:** 10.1101/758060

**Authors:** Arabinda Behera, Gaurav Kumar, Anirban Sain

## Abstract

A semi-rigid filament confined in a soft vesicle of similar size can mutually deform each other. An important example from biological context is Hemoglobin-S (HbS) fibers which polymerize inside red blood cell (RBC). The fibers deform the healthy RBC into sickle-like shape causing difficulty in blood flow through capillaries. Using an area difference elasticity (ADE) model for RBC and a worm-like chain model for the HbS fibers, confined within RBC, we study the shape deformations at equilibrium. We also consider multiple filaments and find that confinement can generate multipolar RBC shapes and can also promote helical filament conformations. The same model, in different parameter regime, reproduces tubulation for phospholipid vesicles, as seen in experiments, when microtubules are confined in the vesicle. We conclude that with a decrease in the surface area to volume ratio, and membrane rigidity, the vesicle prefers tubulation over sickling. Our simulations can access various non-axisymmetric shapes, which have been observed experimentally, both in the context of sickle RBC and phospholipid vesicles, but have so far remained beyond the scope of variational methods.

---

Shape of animal cells and their functions are interrelated. Abnormal cell shape can lead to pathological situations. For example, healthy, red blood cells (RBC) smoothly flow through our circulatory system, where as deformed, sickle shaped RBC cannot pass through capillaries, the thinnest of the blood vessels in our body, giving rise to sickle cell disease. The deformation is caused by semi-rigid Hemoglobin fibers that push the RBC membrane from inside. The fibers grow due to abnormal polymerization of Hemoglobin-S (HbS) molecules and the polymerization is triggered by oxygen deficiency in the blood of patients who have certain genetic defects. Fig.1A shows variety of deformed RBC shapes that are found during sickling and here we ask how the fiber-membrane interaction gives rise to such shapes.

**FIG. 1:**
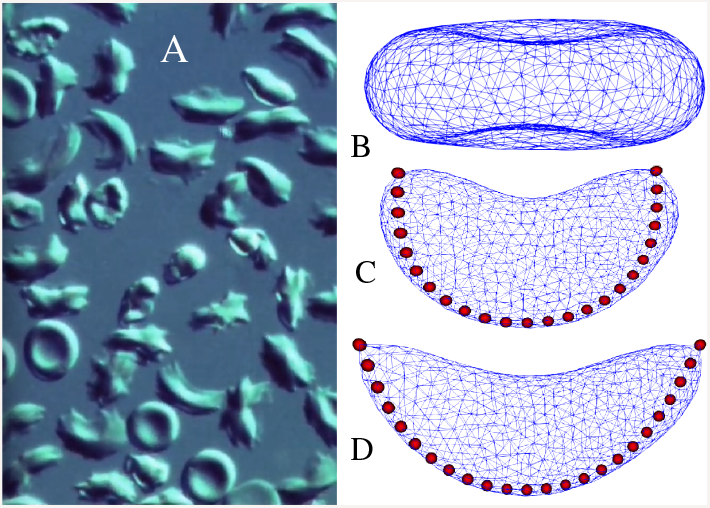
(A) Sickling of RBC observed under microscope (adapted from youtube video, Developmental Biology Film Series (episode 31), UMass Amherst Libraries). The circular shapes with a cavity are the normal RBC; among the deformed RBCs some have sickle shapes and the rest have more complicated multipolar structures. B) Shows the normal biconcave RBC (discocyte) from our simulation, at parameters *κ* = 67.6*k*_*B*_*T* [18], Δ*a*_0_ = 1.03, 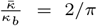, at a fixed reduced volume *ν* = 0.72 and fixed reduced area *a* = 1. (C) and (D) show deformed RBCs with a confined WLC chain indside. The dimensionaless parameter *k*_*B*_*Tl*_*p*_/*k*_*b*_*ρ*_*s*_ = 1.7 and 3.4, for (C) and (D) respectively, where only *l*_*p*_ is doubled from 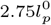 to 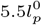.

Each HbS fiber, which we will henceforth refer to as filament, is made of seven double strands of HbS molecules [1] interwined into a twisted structure. Cryo-electron microscopy (CryoEM) has revealed [2] that, inside RBC, HbS fibers occur both as single filament as well as in a bundle form. Possibility of heterogeneous nucleation on these bundles resulting in branched structures has also been considered [3] in the literature. In addition, the organization of the cytoskeleton underlying the RBC membrane may also change [4] during sickling. Here we will focus on elastic deformations of RBC purely due to stiff filaments or filament bundles, confined inside.

A related problem, namely, in vitro polymerization of microtubules (MT) inside phospholipid vesicle and resulting deformations of the vesicle, offers useful insight [5–7]. This is despite the obvious dissimilarity that phospholipid vesicles when pushed from inside by MT filaments, develop tubular protrusions, which is not seen in RBC sickling. We will show that the same model can be used for both the problems, albeit in a different region of the parameter space.

Analytical calculations reported in the context of vesicle tubulation [6, 8, 9] are variational in nature: the structure of the deformed vesicle was assumed to be that of two cylindrical membrane tubes attached to a central ellipsoidal vesicle. The structural parameters like the major and minor axes of the vesicle and the length and radius of the tubes are determined by minimizing the total free energy of the system, often subject to fixed area and fixed volume constraints. This leaves out many of the non-axisymmetric vesicle deformation and almost all sickle conformations of RBC seen in nature (see Fig.1). Numerical free-energy minimization schemes like, Monte-Carlo (MC), equilibrium Molecular dynamics (MD) have been used to study both axisymmetric [10, 11] and non-axisymmetric [12] vesicle deformations. While most of these equilibrium models dealt with filaments with fixed length confined in a vesicle, Ref[13] has used stochastic dynamics to include force-dependent poly/depolymerization mechanisms operating at the filament tips.

## Model

RBC shape morphology, in particular, the main sequence of RBC shapes: stomatocyte, discocyte, and echinocyte have been studied in great detail. Other than its specific area to volume ratio, the membrane properties of RBC has also turned out to be an important determinant of its shape. Using the area difference elasticity(ADE) model Ref[14–16] have shown that by changing a single parameter, namely, the intrinsic area difference Δ*A*_0_, between the two leaflets of the membrane, one can access all the shapes in the main sequence. Here we work with the same RBC parameters that produce discocyte, the healthy biconcave precursors of the sickle RBC as seen under microscopes (see Fig.1-A). With a semi-rigid HbS filament confined inside an RBC, the total free energy of this composite system consists of bending energies of the membrane and the filament, and area difference elasticity of the RBC.

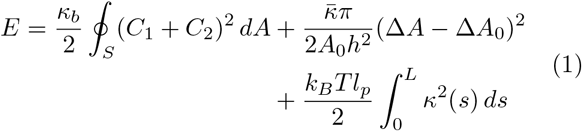

The first term is the Helfrich energy, where *C*_1_ and *C*_2_ are the local principal membrane curvatures. The second term is the non-local bending energy, which arises due to the area difference between the lipid bilayers[17]. Here, Δ*A* = *h* ∮(*C*_1_ + *C*_2_)*dA* is the area difference between the inner and outer phospholipid monolayers, with separation *h* between their neutral planes. The third term represents bending energy of the filament, represented by a worm-like-chain (WLC) model: 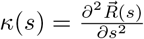 is the local curvature of the filament and *l*_*p*_ is its persistence length, measure of the filament stiffness.

Here we have ignored the effects of cytoskeletal spectrin network beneath the membrane, which gives enhanced elasticity to RBC. Without spectrin budding occurs in ADE model at high Δ*A*_0_ values as spectrin prohibits large stretching and shear deformations that occur at the neck of the buds. Spectrin network is also important for RBC shape fluctuation and its low-frequency dynamic response when RBC undergo large deformations [18]. Since here we restrict ourselves to relatively small values of Δ*A*_0_, same as that of discocyte, ignoring spectrin could be a reasonable approximation. Indeed we reproduce many of the irregular sickle RBC shapes seen under the microscope.

The dimensional-less form of free energy reads

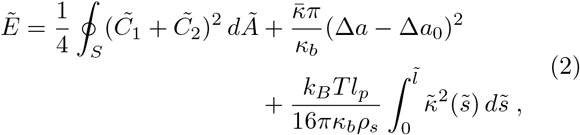

where energy is scaled by 8*πκ*_*b*_, bending energy of a sphere independent of its absolute size, and lengths are scaled by *ρ*_*s*_ which is the radius of the spherical vesicle with the same area as that of RBC. Therefore the reduced area is 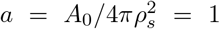 and reduced volume 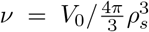. Further, 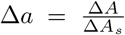 and Δ*a*_0_ = Δ*A*_0_/Δ*A*_*s*_, where Δ*A*_*s*_ = 8*πhρ*_*s*_ is the area difference of a sphere. The filament length and bending energy are also scaled by *ρ*_*s*_ and 8*πκ*_*b*_, respectively. As noted earlier [8] the important parameters in this model are the dimensionless ratios (*k*_*B*_*Tl*_*p*_/*κ*_*b*_*ρ*_*s*_), Δ*a*_0_ and the reduced filament length 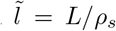. Note, that for a free WLC the only relevant parameter is *l*_*p*_/*L*, but here in the confined case, all lengths have been rescaled by *ρ*_*s*_ which is connected to the area of the RBC. While studying RBC deformations, we vary only *L* and *l*_*p*_, i.e., properties of the filament, and retain all the parameters involving RBC same as that of a healthy bi-concave discocyte. Note that we have not included any intrinsic curvature *C*_0_ in the membrane bending energy. As shown in Ref[16, 19], such an explicit intrinsic curvature would only renormalise the intrinsic area difference elasticity to 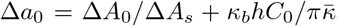. Conversely, the area difference elasticity is also equivalent to an intrinsic curvature.

The vesicle was represented by a discrete, triangulated, closed network with *N* vertices, *N*_*L*_ = 3(*N* − 2) links and *N_T_* = 2(*N* − 2) triangles [20], which satisfy the Euler criteria *χ* = *N* − *N_L_* + *N_T_* = 2 for a convex surface. The HbS filament was represented by a Kratky-Porod chain [21] which is a discrete version of WLC. The chain consists of *M* linearly connected links, with its energy given by

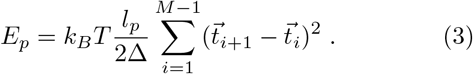

Here 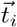’s are link vectors and Δ is the link length such that the filament length is *L* = *M*Δ. We verified that our results are robust with respect to the discretization *M*. Each of the vertices, on the triangulated surface, were allowed to move in order to minimize free energy. Similar in spirit with previous numerical works [10, 12, 13] we confined the filament to the interior of the RBC by using short-range repulsive 2-body potential between the monomers of the filament and the vertices of the membrane. We used hard-sphere potential: 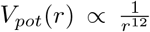, where *r* is separation distance. Note that the internal pushing forces on the membrane due to the confined filament is not constant [22], but depends on the particular filament conformation inside.

In all our simulations a healthy biconcave RBC (discocyte) was taken as the initial shape. The two ends of the filament were attached to the pair of vertices which were farthest in the discocyte shape. This was possible because the filament length considered was always larger than the RBC diameter. To simulate fluidity of the membrane, link-flip moves were carried out. In this move, a link connecting a pair of vertices was picked at random. Erasing this link a new link was formed between another pair of vertices which were common neighbours to previously linked pair [20] but were not linked previously. The acceptance of such a link-flip move was subjected to the minimization of free energy. As a result, all the vertices (including the two which were attached to the filament ends) could diffuse on the triangulated network. Successful Monte-Carlo moves of the vertices were also restricted to the constraints that, a) length of any link must be within an allowed range [*l*_*min*_, *l*_*max*_], and b) the angle between any two neighboring faces (triangles) was below a certain cut-off. This ensured that the network obey self-avoidance criteria and also would not fold onto itself. Even the link flips were subjected to these constraints.

## Results

Our Monte-Carlo simulation produces the most common sickle RBC shapes observed under a microscope, see Fig.1A. But these shapes are obtained (Fig.1D) when the persistence length *l*_*p*_ is taken to be about six times higher compared to that of a single HbS fiber 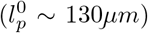 [23]. This is consistent with, a) CryoEM studies [2] which detected bundles in frozen, hydrated, sickle HbS fibers, and b) in vitro measurements of *l*_*p*_ [24] which reported a range of persistence lengths varying about 50 times, suggesting up to seven fibers could be bundled together (as 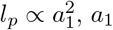 being the cross-sectional area of a fiber). The area and volume of the initial biconcave discocyte were taken to be 140*μm*^2^ and 100*μm*^3^ [14], respectively. The length of the semi-rigid filament was chosen to be longer than the initial discocyte diameter and its initial configuration was a random coil. Constant area and volume were maintained during each MC run. Sickling increased for higher *l*_*p*_. Fig.1C and D are produced for 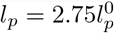 and 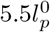, respectively.

In Fig.2, we show deformations due to change in the dimensionless filament length 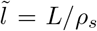. Larger *L* leads to higher bending, and superficially this appears to be equivalent to lowering *l*_*p*_ as in Fig.1. But note that unlike for a free WLC where *l*_*p*_/*L* is the only relevant parameter for determining filament conformation, here in the presence of a confinement length scale *ρ*_*s*_ we can have two very different filament conformations at same *l*_*p*_/*L*. In Fig 2B,D we show two such conformations which have the same *l*_*p*_/*L*, but in Fig2D both *l*_*p*_ and *L* are 1.5 times of that in Fig2B.

**FIG. 2:**
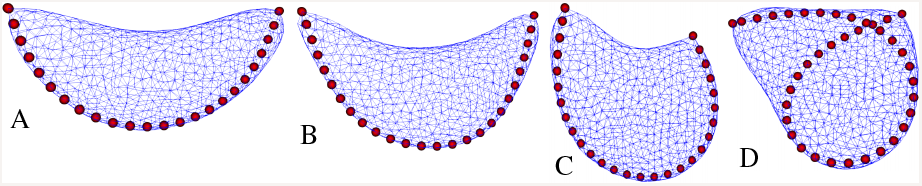
Sickle RBC shapes A,B and C, for increasing filament lengths *L* = 46, 50, 54 in arbitrary units. The parameters *k*_*B*_*Tl*_*p*_/*k*_*b*_*ρ*_*s*_ and Δ*a*_0_ are same as in Fig.1-D. In (D) both *l*_*p*_ and *L* were made 1.5 times of that in (B). As a result the filament collapsed instead of elongating the RBC. Fixed volume and area, as in Fig.1 were maintained.

We also note that up to a critical length *L*_*c*_, the HbS filaments, although already buckled, continue to push the membrane hard creating pointed ends. But beyond *L*_*c*_ the filament collapses to circular arcs and folds back on itself, and the pointed corners of the vesicle become rounded. Quantitatively, the end-to-end distance of the filament undergoes a jump. Similar observation has also been made for MT filaments growing inside phospholipid vesicles. This effect has been explained in Ref[25] using a phenomenological theory and the maximum longitudinal force that a bent filament can sustain before the collapse has been found numerically by solving a transcendental equation. In the light of this discussion, we note that most of the sickled RBC in Fig.1 have many pointed ends indicating that that most of the HbS filaments inside do not cross this critical *L*_*c*_. This could be because, a) the filament nucleation rate is sufficiently high to generate many filaments that grow and reach the boundaries simultaneously, and b) long filaments are intrinsically unstable and the system favours branched structures resulting from heterogeneous nucleation.

The model which we used for RBC, can also account for the shapes and shape transition of lipid vesicles when MT or MT-bundles polymerize inside the vesicle [5, 6]. MT has a high persistence length, of order 6*mm* [5, 26], whereas phospholipid vesicle has relatively low bending rigidity *κ*_*b*_ ≈ 10*k*_*B*_*T*. This makes the nondimensional parameter *k*_*B*_*Tl*_*p*_/*κ*_*b*_*ρ*_*s*_ ≈ 100, about twenty times larger than that of RBC. Given that lipid vesicles without MT inside are nearly spherical at room temperature, we choose *ν* = 0.9 and Δ*a*_0_ ≈ 2. Fig.3 shows tube formation and subsequent bending of the MT filament at higher length. However, our simulation always produces one tube in contrast to two tubes that were reported in Ref[5, 6]. We believe that two tubes occur due to nonequilibrium effects resulting from dynamic polymerization. Ref[12] (and references therein) has shown that the two tubes state is a metastable minima compared to the single tube state. Qualitatively, when the total length of two tubes is same as that of a single tube (of the same radius), the energy difference should be the cost of one kink at the base of the tube. We note that in variational calculations [6, 8, 9] the two tube conformation was assumed apriori. Experiments in Ref[6] with MT filaments showed a reentrant transition for the vesicle shape: *tube → collapsed → tube*, as the MT bundle grew in length. We recover this transition which is shown in Fig.3. In Ref[5] similar *tube → collapsed* transition was reported by increasing the membrane tension (*σ*) by tuning aspiration pressure. Within our simulation, this is equivalent to increasing 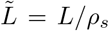, where 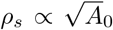. In the constant surface tension ensemble increasing *σ* amounts to decreasing the area *A*_0_. In our constant area ensemble decreasing *A*_0_ is equivalent to increasing length *L* of the MT.

**FIG. 3:**
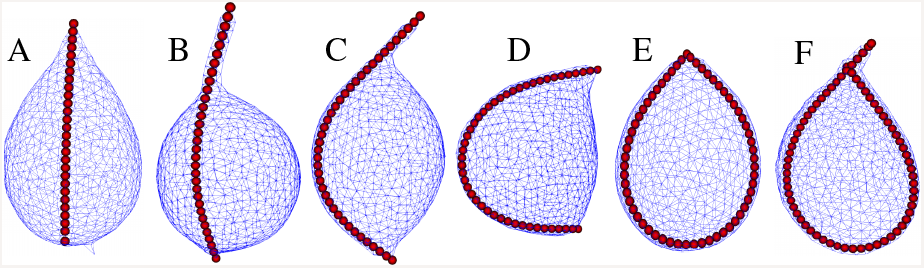
Stable shapes of deformed phospholipid vesicles upon increase in MT length. From A through F, *L* = 30, 32, 40, 48, 60, 69, in arbitrary units. Other parameters are fixed at *k*_*B*_*Tl*_*p*_/*k*_*b*_*ρ*_*s*_ = 111, with *κ*_*b*_ = 10*k*_*B*_*T*, 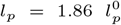 (when 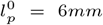), reduced volume *ν* = 0.92 (i.e., nearly sphere) and Δ*a*_0_ = 2.1. Bending of initially straight tube (A → *B* → *C*), shrinkage of tubes (C → D) and collapse of filament into closed conformation (D → E) and retubulation (E → F) can be seen. The filament collapse transition and generation of tennis racket type of shape were experimentally reported in Ref [5] and [6], respectively. The filament appears continuous here due to relatively finer discretization, compared to other figures.

As is manifest in Fig.1A HbS filaments can generate a variety of shapes other than sickle shapes. Some of these have multiple protrusions and may be the result of simultaneous polymerization of more than one filament. There could also be branched HbS filament structures. The amount of HbS molecules vary across patients giving rise to variability in the number of filaments that can form in a sickle RBC. In Fig.4 we examine deformation of RBC shape due to two HbS filaments. Often, starting from the same initial configuration, we get different metastable shapes, with 2,3 and 4 poles, which show that the kinetics of the deformation is important in locking the deformations into one shape or other.

**FIG. 4:**
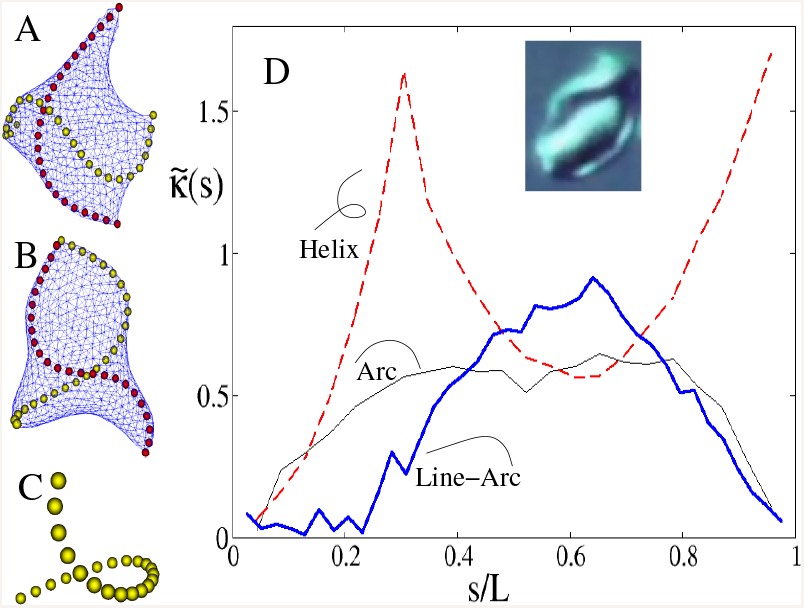
Multipolar RBC shapes due to two filaments : A and B represent metastable minima with B’s energy 5% lesser than A. They had same initial conditions, with the filaments arranged perpendicular to each other along the diameters of a healthy RBC. Often the filaments are helical. One of the filaments in B is separately shown in C; its helical conformation is clearly visible. Parameters are Δ*a*_0_ = 1.03, *k*_*B*_*Tl*_*p*_/*κ*_*b*_*ρ*_*s*_ = 17 (with 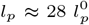), 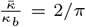, *ν* = 0.72, *a* = 1. D) Nonuniform local curvature 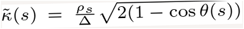, where *θ* is the angle between successive tangent vectors, versus reduced contour length *s/L*, for qualitatively different shapes (see legends). The bold line (line-arc) is for the MT filament in Fig.3C, and the rest two are for HbS filaments in sickle RBC. Only the planar arc conformation (from Fig.1D) shows a significant plateau, indicative of uniform curvature. All the helical conformations which we examined (dashed line is for C), show sharp kinks in curvature. The MT bundle (line-arc) shows zero curvature inside the tube, but nonuniform curvature within the ellipsoidal portion. The inset shows a plausible case of helical HbS filament inside a sickle RBC (source: same as Fig.1A)

Semiflexible filaments, described by WLC model, do not form helix by themselves when confined between two walls. Such 1D confinement leads to planar buckling. As the walls are brought closer, the buckling amplitude grows into nonlinear regime. Various authors [27, 28] have obtained helical low-energy structures in discrete WLC model by incorporating beyond nearest-neighbor (semi-long range) attarction between non-bonded monomers. In the context of proteins, maximum compactness criteria can also lead to helix [29]. In our present case of confined HbS fibers, 3D confinement effect gives rise to helix, see Fig.4A-C. In Fig.4D we highlight the curvature distribution along the contour of the filaments for various conformations discussed so far. Interestingly, the curvatures are highly non-uniform, making it difficult to predict such conformations analytically.

In summary we showed that, a) inclusion of the semi-rigid filaments into normal RBC is sufficeint to reproduce the basic sickle shape morphology. b) It turns out that a single HbS fiber is not stiff enough to reproduce the sickle RBC shapes with pointed corners, and we predict that HbS fiber bundles must be involved. c) Our model qualitatively reproduces the tube collapse transition and tennis racket type shape, seen in experiments on phospholipid vesicles with MT filament bundles inside. We showed that even in sickled RBC similar transitions are theoretically possible. Absence of the same in diseased blood samples points to the possibility of heterogeneous nucleation and/or some other instability for length control in HbS filaments. d) We also predict helical HbS conformations arising from 3D confinement effect. Our MC simulation cannot capture the nonequilibrium polymerization kinetics, which we think is responsible for, a) giving rise to the metastable two-tubes state, in the context of MT driven vesicle tubulation, and b) multiple pointed corners in sickle RBC due to heterogeneous nucleation and/or simultaneous growth of multiple filaments. Clearly, any chemical means that could reduce the persistence length of HbS filaments, by suppressing bundling, can oppose sickling.

We thank Prof. Debjani Paul’s group (IIT Bombay) for introducing us to the phenomenology of sickle cell disease and for many useful discussions. We acknowledge Space-Time-2 supercomputing facility at IIT Bombay. AB and GD would like to thank UGC-India and IRCC-IIT Bombay, respectively for financial support.

